# Schizophrenia polygenic risk scores, urbanicity and treatment-resistant schizophrenia

**DOI:** 10.1101/501288

**Authors:** Christiane Gasse, Theresa Wimberley, Yungpeng Wang, Henriette Thisted Horsdal

**Author notes:** Corresponding author: Christiane Gasse, PhD, Dip Epid (McGill), Forskningsfarmaceut, Mobil: 51191476, Aarhus Universitetshospital Psykiatry, Department of Depression and Anxiety/Psychosis Research Unit, Palle Juul-Jensens Boulevard 175 ■ DK-8200 Aarhus N, Denmark.

## Abstract

**Introduction:** To investigate the impact of a polygenic risk score for schizophrenia (PRS-SZ) and urbanicity on the risk of treatment-resistant schizophrenia in people with a diagnosis of schizophrenia and to evaluate the association between PRS-SZ and TRS across areas of urbanicity.

**Methods:** Cohort study of people born after 1981 with a first time diagnosis of schizophrenia between 1996 and 2012 using Danish population registry data. Through linkage to genome-wide data, we calculated PRS-SZ based on a Psychiatric Genomics Consortium meta-analysis. We assessed urbanicity at birth (capital, provincial and rural areas). TRS was defined using prescription and hospital data. Performing Cox regression analysis, we calculated hazard rate ratios (HRs) and 95% confidence intervals (CI).

**Results:** Among 4,475 people with schizophrenia, we identified 593 (13.3%) with TRS during 17 558 person years of follow-up. The adjusted HR for TRS associated with 1 standard deviation (SD) increase in the PRS-SZ was 1.11 (95% CI: 1.00–1.24). The adjusted HRs for urbanicity and TRS were 1.20 (95% CI: 0.98–1.47) for provincial areas and 1.19 (95% CI 0.96–1.47) for rural areas compared with the capital area. Across strata of urbanicity, the adjusted HR for TRS was 1.39 (95% CI: 1.14–1.70) in the capital area with 1 SD increase in the PRS-SZ, 0.99 (95% CI 0.84–1.17) in provincial areas, and 1.03 (95% CI: 0.86–1.25) in rural areas.

**Conclusion:** The risk of TRS associated with genetic liability varied across urbanicity areas and was highest in people with schizophrenia who resided in the capital areas at birth.

## 1 INTRODUCTION

Insufficient response to antipsychotic treatment, also termed treatment-resistant schizophrenia (TRS) may affect up to 30% of people with schizophrenia initiating antipsychotic treatment (Meltzer, 1997; Howes et al., 2017). The reasons for TRS are still not well understood (Gillespie et al., 2017;Wimberley et al., 2016b). Among a number of patient and treatment related factors, we have previously shown that living in rural areas at birth was associated with increased risk of TRS (Wimberley et al., 2016b;Wimberley et al., 2016a). Specific genetic factors in relation to TRS have not been identified yet (Gillespie et al., 2017;Lally et al., 2016), though a few studies point towards a weak association between a polygenic risk score for schizophrenia (PRS-SZ) and TRS, indicating higher genetic liability for schizophrenia among people with TRS (Wimberley et al., 2017;Frank et al., 2015). The precision of these genetic studies was though potentially affected by low power of the sample sizes. Another study from Denmark showed that people with higher PRS-SZ tend to be born (and to live) more frequently in urban areas, thereby indicating that people in rural areas showing lower PRSs for schizophrenia (Paksarian et al., 2018). Thus, the lower PRS-SZ but increased risk of TRS in people born or living in rural areas lead us to the hypothesis that urbanicity and PRS-SZ may interact resulting in different estimates of the associations between PRS and TRS in different geographical areas.

Therefore, we investigated here: (1) whether we could replicate and estimate more precisely our previous findings of an association between both a PRS-SZ and urbanicity and TRS using the largest population-based sample of people with schizophrenia; (2) whether levels of PRS-SZ varied in people with and without TRS in different geographical areas.

## 2. METHODS

### 2.1 Study design and data

We performed a prospective cohort study by linking Danish administrative registries that cover the entire Danish population using the unique civil registration number assigned to every person living in Denmark and thereby enabling linkage between registers. We obtained data from the following Danish national registers: The Danish Civil Registration System (Pedersen, 2011) containing records on gender, date of birth, change of address, date of emigration, vital status, and links to family members since 1968. The Danish Psychiatric Central Research Register containing all discharge diagnoses assigned at psychiatric hospitals in Denmark since 1969 (outpatient contacts included since 1995) (Mors et al., 2011). The diagnoses are coded according to International Classification of Disease (8th revision) (ICD-8) until the end of 1993, and 10th revision (ICD-10) thereafter. The Danish National Prescription Registry containing the date, kind of drug and volume of redeemed prescriptions at community pharmacies, registered since 1995 (Pottegard et al., 2017). The Danish Newborn Screening Biobank (Norgaard-Pedersen and Hougaard, 2007) containing genetic data extracted from dried blood spots collected in the first days after birth of nearly all infants born in Denmark since 1981.

### 2.2 Study population

Among individuals born after May 1, 1981 in Denmark, we identified people with a first diagnosis of schizophrenia as in- or out-patient or emergency room contact at a psychiatric hospital (ICD-10: F20) between January 1, 1996 and December 31, 2012. Thus, individuals with a ICD-8 diagnosis (295.x9 excluding 295.79) or a ICD-10 diagnosis of schizophrenia recorded before 1996 were not included. Furthermore, we included individuals only if they were 10 years or older at the first registered diagnosis of schizophrenia and alive at discharge, and had a known mother identified by linkage to the Danish Civil Registration System (Pedersen, 2011). To the identified individuals, we linked prescription data from the Danish National Prescription Registry and retrieved information on antipsychotic prescriptions (Anatomical Therapeutic Chemical (ATC) Classification system code N05A) (10). We excluded 19 people with a clozapine prescription before their first registered diagnosis of schizophrenia. The final study population consisted of the individuals with a DNA sample available from the Danish Newborn Screening Biobank (Norgaard-Pedersen and Hougaard, 2007).

### 2.3 Outcome measure

#### 2.3.1 Treatment-resistant schizophrenia

Our measure of TRS was based on hospital admission data from the Danish Psychiatric Central Research Register (Mors et al., 2011) and the Danish National Prescription Registry (Pottegard et al., 2017) regarding antipsychotic prescription redemptions. We defined TRS as the first occurrence of either clozapine initiation or hospitalization due to schizophrenia during antipsychotic treatment within 18 months after at least two periods of different antipsychotic monotherapies lasting at least 6 weeks each. This treatment-based definition, used in recent published studies, builds on international and Danish treatment guidelines and the Kane criteria (Wimberley et al., 2017;Wimberley et al., 2016b). We recently validated this algorithm against clinical records in UK and found a positive predictive value of 63.6%, and a sensitivity of 62.2% compared with a positive predictive value of 78.3% and a sensitivity of 40% for identifying TRS based on clozapine prescriptions only (Ajnakina et al., 2018).

### 2.4 Exposure measures

#### 2.4.1 The polygenic risk score

DNA from the genetic material obtained from dried blood spot samples of the study population was whole-genome amplified in triplicate using the Qiagen REPLI-g mini kit [Qiagen, Hilden, Germany]. The three separate reactions were pooled (Hollegaard et al., 2009;Hollegaard et al., 2011) and genotyped with either Illumina Human 610-Quad BeadChip array, Illumina HumanCoreExome beadchip or Illumina Infinium PsychArray-24-v1.1 BeadChip (Illumina, San Diego, CA) (Pedersen et al., 2018;Agerbo et al., 2012;Meier et al., 2016).

We calculated PRS-SZ based on the summary statistics (effect allele, effect size) derived from the Psychiatric Genomics Consortium (PGC) discovery sample (2014). The discovery sample comprised 34,600 schizophrenia cases and 45,986 controls from the PGC Genome-Wide Association Study (GWAS) meta-analysis for schizophrenia, excluding the Danish cases (2014). We selected single-nucleotide polymorphisms (SNPs) associated at a p-value threshold of 0.05 or lower. This threshold was chosen in accordance with other studies including polygenic risk scores (Wimberley et al., 2017;Agerbo et al., 2015) and is considered appropriate to achieve a balance between the number of false-positive and true-positive risk alleles (Wray et al., 2014). In our target sample, our independent study population, we calculated a polygenic risk score for each individual as the weighted sum of risk alleles at the selected SNPs with the weight being the effect estimates in the discovery sample. The calculated PRS-SZ, that was approximately normally distributed (Dudbridge, 2013), was standardized by subtracting the mean divided by the square root of the sample variance.

#### 2.4.2 Urbanicity

We obtained information on geographical area at birth from the Civil Registration System (8). Geographical areas were categorized into three levels according to the degree of urbanization: capital area (capital and suburbs of the capital), provincial areas (>10,000 inhabitants), and rural areas (≤10,000 inhabitants).

### 2.5 Covariates

We obtained information on age at first schizophrenia diagnosis and gender from the Civil Registration System (Pedersen, 2011). The year of diagnosis of schizophrenia was obtained from the date of the first diagnosis of schizophrenia recorded in the Danish Psychiatric Central Research Register (Mors et al., 2011). Because the samples were genotyped using three different types of arrays as described above, we accounted for the sampling frames in the statistical analyses by adjusting for the different genotyping arrays. Ancestry was assessed by the first 10 genomic principal components (Price et al., 2006).

### 2.6 Statistical analysis

We followed individuals from the date of their first diagnosis for schizophrenia until TRS, emigration, death or end of study period (June 30, 2013). The mean PRS-SZ for each level of urbanicity was estimated and adjusted for age at diagnosis, sex, year of diagnosis, the three types of genotyping arrays, and the 10 first genomic principal components. We performed Cox regression analysis and calculated hazard rate ratios (HRs) for TRS in relation to a PRS-SZ and urbanicity. We have previously shown that females are at increased risk of TRS in Denmark and allowed therefore different baseline hazards for males and females in the models (Wimberley et al., 2016b). The HRs were mutually adjusted for PRS-SZ and urbanicity and additionally for age at diagnosis, year of diagnosis, the three types of genotyping arrays, and the 10 first genomic principal components. We tested for an interaction between PRS-SZ and urbanicity. In case of an interaction, we planned to evaluate further the association between PRS-SZ and TRS according to levels of urbanicity. To assess the robustness of our results we performed a sensitivity analysis where we restricted the TRS definition to people who redeemed at least one prescription for clozapine, which has been shown to have a better positive predictive value though lower sensitivity compared with the algorithm derived TRS status (Ajnakina et al., 2018).

All statistical analyses were performed using STATA 13.1 and SAS 9.4.

## 3 RESULTS

We included 4475 people with a first registered diagnosis of schizophrenia between 1996 and 2012 and available genetic information. Characteristics of the study population are shown in Table 1. The median age at first schizophrenia diagnosis was 20.6 years (interquartile range 18.4 years to 23.1 years) and 43% were female. Slightly more than a third lived in the provincial areas at birth, and the third sampling frame contributed 61% of the cases (Table 1). The study population was followed for 17 558 person years (Table 2). During follow up, 61 people died, and 65 were censored due to emigration. 593 (13.3%) fulfilled the criteria for TRS during follow-up at a rate of 3.38 cases of TRS per 100 person years.

**Table 1:** Baseline characteristics at first schizophrenia diagnosis, n=4,475.

| Baseline characteristics | Total |
| --- | --- |
| N (%) | 4,475 (100) |
| Age, median (inter-quartile range) | 20.6 (18.4-23.1) |
| Females, n (%) | 1,944 (43.4) |
| Geographical area at birth, n (%) |  |
| Capital region | 1,391 (31.1) |
| Provincial region | 1,702 (38.0) |
| Rural region | 1,382 (30.9) |
| Genotyping array/sampling frame*, n (%) |  |
| 1 | 873 (19.5) |
| 2 | 885 (19.8) |
| 3 | 2,717 (60.7) |
| Year of diagnosis, n (%) |  |
| 1996-2000 | 113 (2.5) |
| 2001-2004 | 495 (11.1) |
| 2005-2008 | 1,345 (30.1) |
| 2009-2012 | 2,522 (56.3) |
| PRS, mean (SD) | 2.57x10<sup>-10</sup> |

\*Genotyping arrays used in different sampling frames: (1) Illumina Human 610-Quad BeadChip array; (2) Illumina HumanCoreExome beadchip; (3) Illumina Infinium PsychArray-24-v1.1 BeadChip (Illumina, San Diego, CA)

### 3.1 Associations between PRS, urbanicity and TRS

Crude and adjusted HRs of the associations between PRS-SZ, urbanicity and TRS are displayed in Table 2. The adjusted HR of the association between 1 SD increase in the PRS-SZ and TRS was 1.11 (95% CI: 1.00–1.24). The HRs increased with the PRS-SZ from the lowest to the highest quartile of the PRS-SZ indicating an upward trend. The adjusted HR for the highest versus the lowest quartile was 1.36 (95% CI: 1.06–1.73).

**Table 2:**
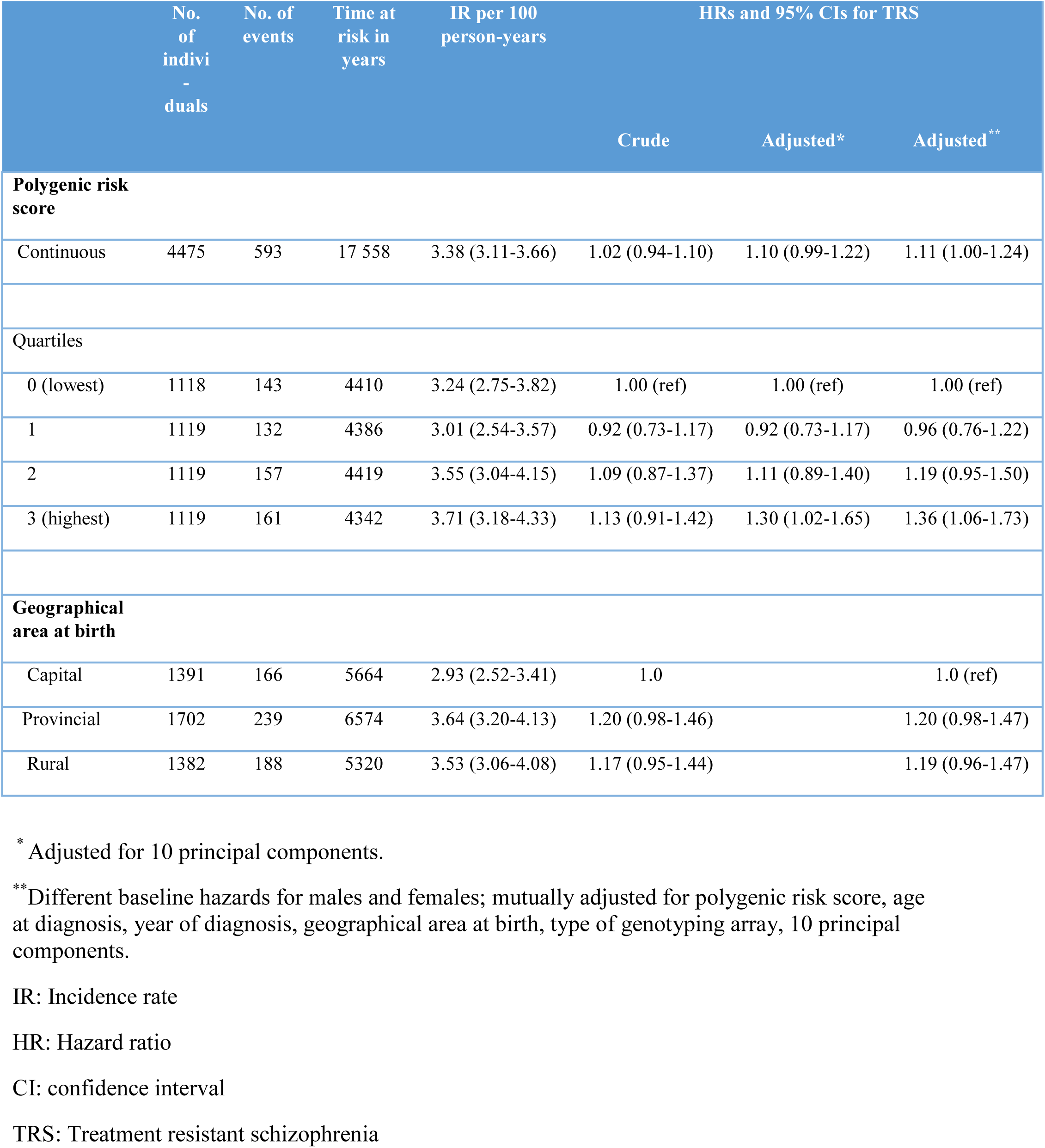
Cox regression analysis of **treatment-resistant schizophrenia** during follow up.

The adjusted HRs of the association between urbanicity at birth and TRS were 1.20 (95% CI: 0.98–1.47) for provincial areas and 1.19 (95% CI 0.96–1.47) for rural areas compared with the capital area.

Modelling the interaction between the continuous PRS-SZ and urbanicity by including an interaction term, the adjusted HR for 1 SD increase in PRS-SZ in association with TRS was 1.34 (1.13–1.58). The HRs for the association between urbanicity and TRS were 1.24 (95% CI 1.00–1.52) for provincial areas, and 1.21 (0.97–1.51) for rural areas compared with the capital area. The p-values for the interaction terms between the continuous PRS-SZ and urbanicity were 0.007 for provincial areas, and 0.068 for rural areas, thus indicating a significant interaction.

For further illustration of the interaction between PRS-SZ and urbanicity on the risk of TRS, the mean PRSs-SZ across levels of urbanicity are displayed in Supplementary Figure 1. We found a trend towards higher PRSs-SZ in the capital and provincial areas compared with rural areas. Figure 1 displays the mean PRSs-SZ additionally stratified by TRS status in the three different levels of urbanicity. The mean PRSs-SZ were significantly higher among people with TRS in the capital area (p-value = 0.0005) compared with those with non-TRS, but not statistically different in the other geographical areas (provincial area: p-value=0.9325; rural area: p-value=0.7021). Table 3 shows HRs for the associations between PRS-SZ and TRS stratified by urbanicity levels. In the adjusted model, 1 SD increase in the PRS-SZ increased the risk of TRS by a HR of 1.39 (95% CI: 1.14–1.70) in the capital area. In provincial and rural areas, the adjusted HRs for the associations between PRS-SZ and TRS were around 1 for 1 SD increase in the PRS-SZ. When analyzing the association between the PRS-SZ and TRS in quartiles of the PRS-SZ, within the capital area, we found an increasing trend of the PRS-SZ associated risk of TRS from the second lowest quartile of the PRS-SZ (HR of 1.55 (95% CI: 0.92–2.62) to the highest (HR of 2.44 (95% CI: 1.44–4.13) within the capital area, (Table 3). This trend though did not reach statistical significance indicated by overlapping confidence intervals. In provincial and rural areas, there was also an indication of an increase in risks for TRS from the lowest to the highest quartiles of the PRS-SZ.

**Figure 1.**
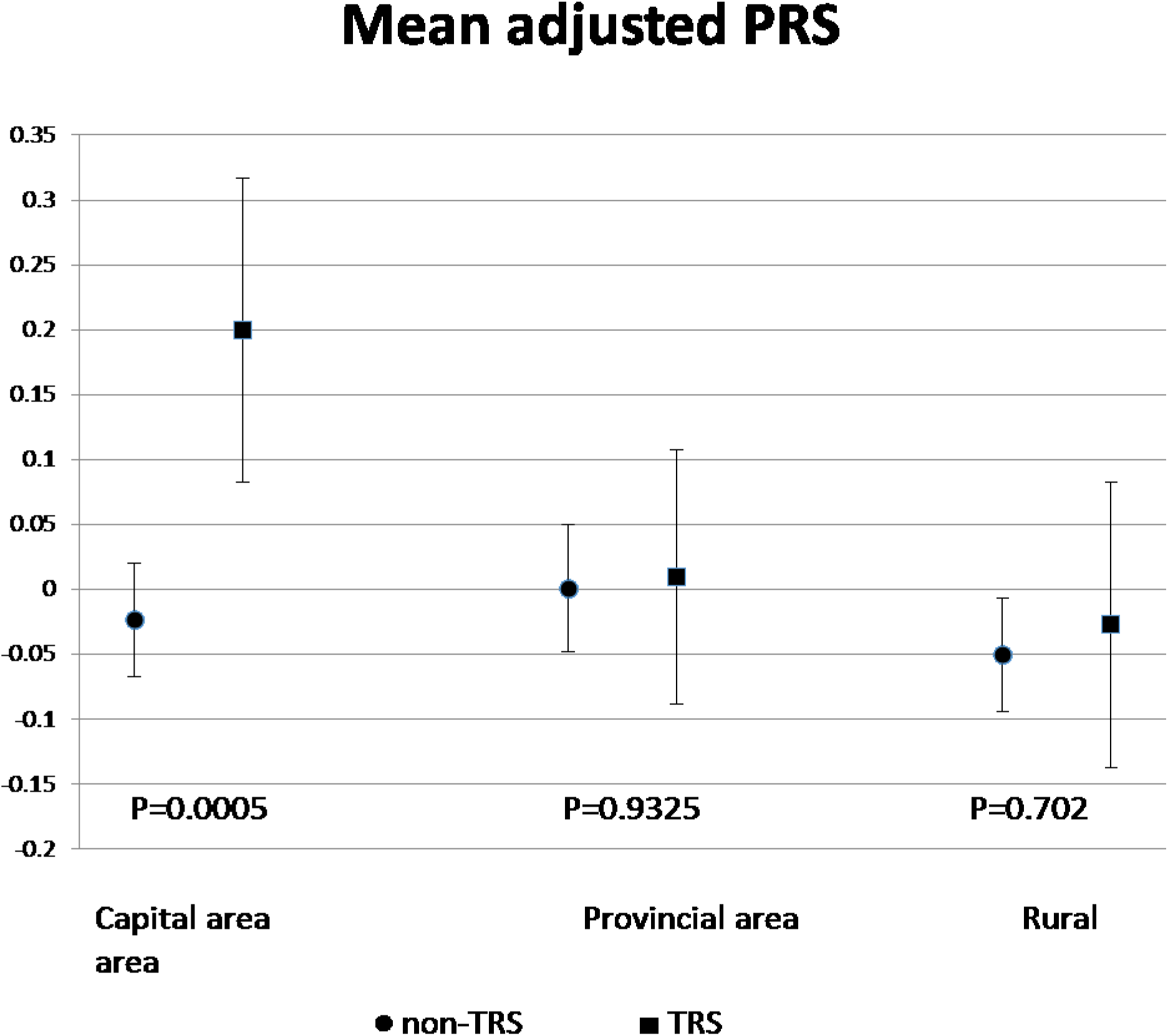
Mean polygenic risk scores for schizophrenia in people with schizophrenia according to geographical area and by TRS status, adjusted for sex, age at diagnosis, year of diagnosis, type of genotyping array, 10 principal components. P-values for the comparison of differences in means of polygenic risk scores.

**Table 3:**
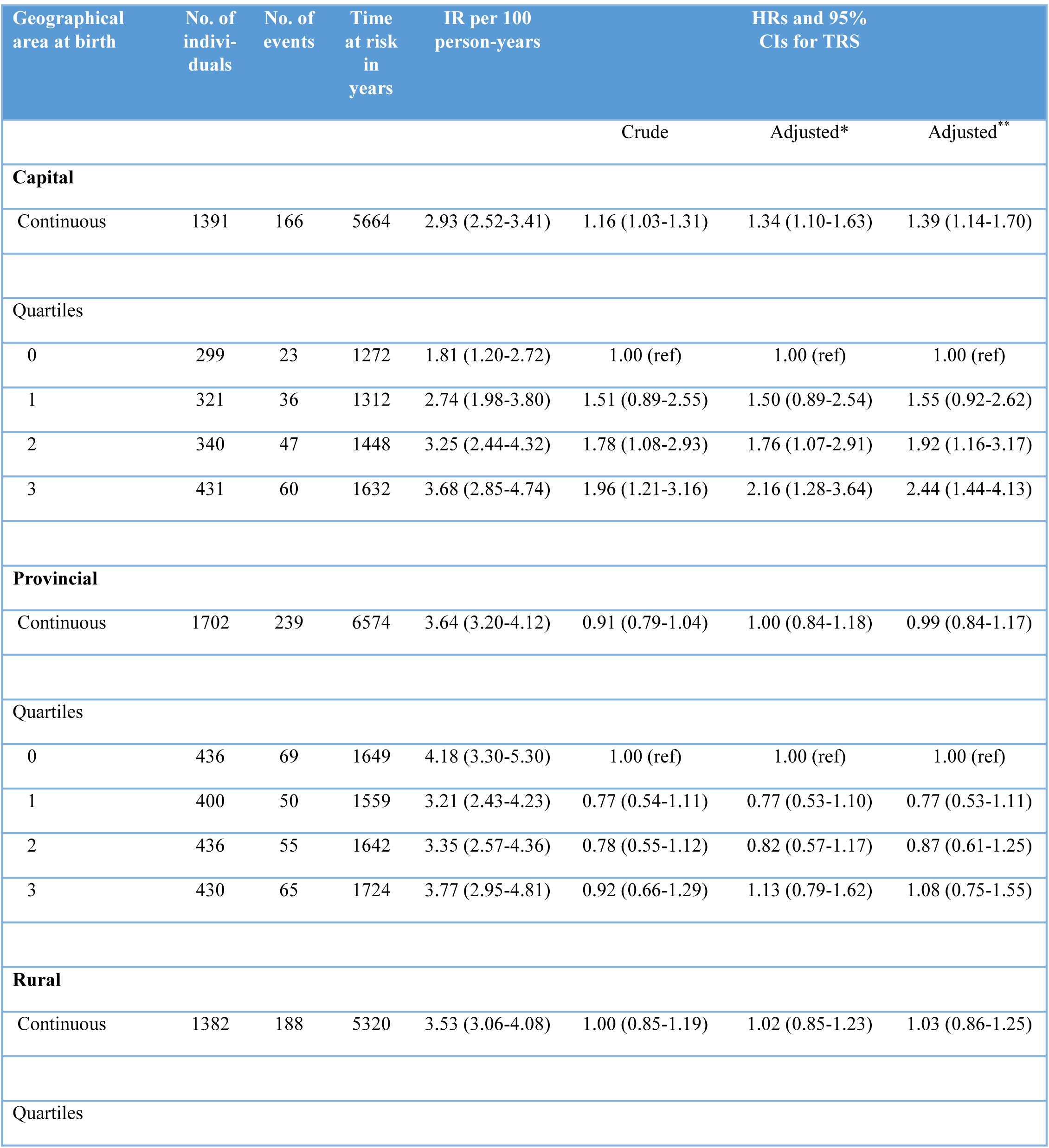
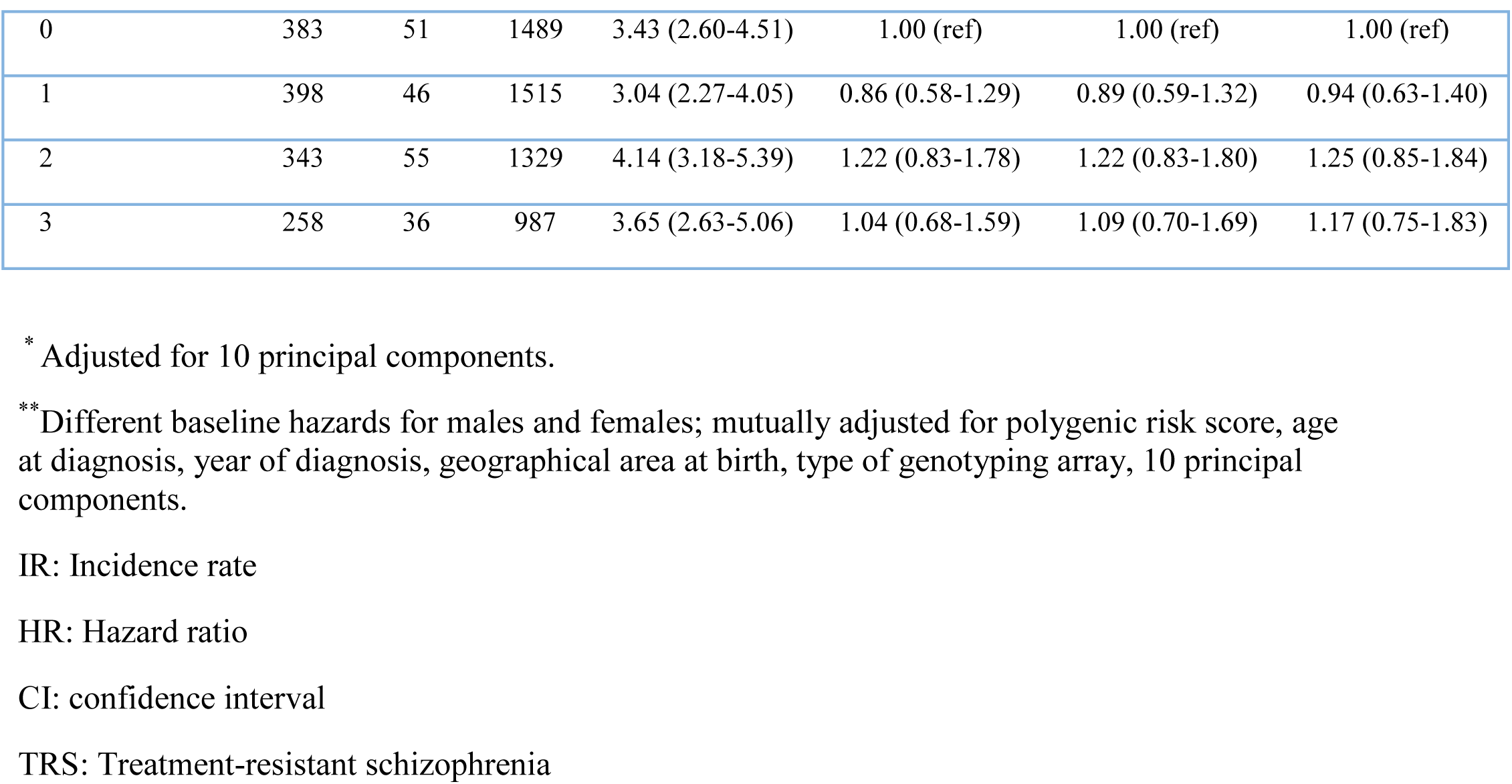
Cox regression analysis of **treatment-resistant schizophrenia** during follow up, stratified by geographical area at birth.

| Geographical area at birth | No. of individuals | No. of events | Time at risk in years | IR per 100 person-years | HRs and 95% CIs for TRS |  |  |
| --- | --- | --- | --- | --- | --- | --- | --- |
|  |  |  |  |  | Crude | Adjusted* | Adjusted** |
| Capital |  |  |  |  |  |  |  |
| Continuous | 1391 | 166 | 5664 | 2.93 (2.52-3.41) | 1.16 (1.03-1.31) | 1.34 (1.10-1.63) | 1.39 (1.14-1.70) |
| Quartiles |  |  |  |  |  |  |  |
| 0 | 299 | 23 | 1272 | 1.81 (1.20-2.72) | 1.00 (ref) | 1.00 (ref) | 1.00 (ref) |
| 1 | 321 | 36 | 1312 | 2.74 (1.98-3.80) | 1.51 (0.89-2.55) | 1.50 (0.89-2.54) | 1.55 (0.92-2.62) |
| 2 | 340 | 47 | 1448 | 3.25 (2.44-4.32) | 1.78 (1.08-2.93) | 1.76 (1.07-2.91) | 1.92 (1.16-3.17) |
| 3 | 431 | 60 | 1632 | 3.68 (2.85-4.74) | 1.96 (1.21-3.16) | 2.16 (1.28-3.64) | 2.44 (1.44-4.13) |
| Provincial |  |  |  |  |  |  |  |
| Continuous | 1702 | 239 | 6574 | 3.64 (3.20-4.12) | 0.91 (0.79-1.04) | 1.00 (0.84-1.18) | 0.99 (0.84-1.17) |
| Quartiles |  |  |  |  |  |  |  |
| 0 | 436 | 69 | 1649 | 4.18 (3.30-5.30) | 1.00 (ref) | 1.00 (ref) | 1.00 (ref) |
| 1 | 400 | 50 | 1559 | 3.21 (2.43-4.23) | 0.77 (0.54-1.11) | 0.77 (0.53-1.10) | 0.77 (0.53-1.11) |
| 2 | 436 | 55 | 1642 | 3.35 (2.57-4.36) | 0.78 (0.55-1.12) | 0.82 (0.57-1.17) | 0.87 (0.61-1.25) |
| 3 | 430 | 65 | 1724 | 3.77 (2.95-4.81) | 0.92 (0.66-1.29) | 1.13 (0.79-1.62) | 1.08 (0.75-1.55) |
| Rural |  |  |  |  |  |  |  |
| Continuous | 1382 | 188 | 5320 | 3.53 (3.06-4.08) | 1.00 (0.85-1.19) | 1.02 (0.85-1.23) | 1.03 (0.86-1.25) |
| Quartiles |  |  |  |  |  |  |  |
| 0 | 383 | 51 | 1489 | 3.43 (2.60-4.51) | 1.00 (ref) | 1.00 (ref) | 1.00 (ref) |
| 1 | 398 | 46 | 1515 | 3.04 (2.27-4.05) | 0.86 (0.58-1.29) | 0.89 (0.59-1.32) | 0.94 (0.63-1.40) |
| 2 | 343 | 55 | 1329 | 4.14 (3.18-5.39) | 1.22 (0.83-1.78) | 1.22 (0.83-1.80) | 1.25 (0.85-1.84) |
| 3 | 258 | 36 | 987 | 3.65 (2.63-5.06) | 1.04 (0.68-1.59) | 1.09 (0.70-1.69) | 1.17 (0.75-1.83) |

\* Adjusted for 10 principal components.
\*\* Different baseline hazards for males and females; mutually adjusted for polygenic risk score, age at diagnosis, year of diagnosis, geographical area at birth, type of genotyping array, 10 principal components.
IR: Incidence rate
HR: Hazard ratio
CI: confidence interval
TRS: Treatment-resistant schizophrenia

### 3.2 Sensitivity analyses

In the sensitivity analyses, we compared people initiating clozapine, as an alternative definition of TRS, versus non-clozapine users, thereby not excluding people classified as TRS using the algorithm based TRS definition. We found similar estimates as for the broader TRS definition, though providing statistically insignificant results, potentially due to missing power and TRS case misclassification among controls of non-clozapine users (Supplementary Table 1). The HRs of the association between urbanicity at birth and clozapine were up to 38% increased in both provincial and rural areas compared with the capital area. The p-values for the interaction between the continuous PRS and urbanicity (3 levels) was p=0.08 for provincial areas; and p=0.5 for rural areas.

## 4 DISCUSSION

In this currently largest population-based sample of people with schizophrenia and TRS, we found a weak effect of a 10% increased genetic liability for schizophrenia, as measured by a polygenic risk score, in association with TRS. Comparing the highest quartile with the lowest of the PRS-SZ, thus indicating higher load of genetic risk in the highest quartile, the risk for TRS was 36% increased. Results using clozapine as an indicator for treatment resistance corroborated these findings. Our overall finding lies in the range of previous estimates of the association between a PRS-SZ and TRS (Wimberley et al., 2017)(Frank et al., 2015). Thus, the impact of genetic liability for schizophrenia as measured by a PRS-SZ on risk of TRS appears low. We previously found no association between family history of schizophrenia and TRS (Wimberley et al., 2016b).

Regarding urbanicity, we found a 20% increased risk for TRS in provincial and rural areas. This is in line with our previous finding of a HR of 1.19 (95% CI: 1.05–1.34) for rural areas and 1.12 (95% CI: 0.98–1.28) in a larger and slightly older study population with a median age of 28 years (Wimberley et al., 2016a). In the study by Wimberley and colleagues assessing urbanicity at the time of the first schizophrenia diagnosis, the association between rural and provincial areas and TRS was higher at HRs 1.60 (95% CI: 1.43–1.79) and 1.44 (95% CI: 1.31–1.59), respectively (Wimberley et al., 2016a).

We detected an interaction between urbanicity and genetic liability for schizophrenia on the risk of TRS. We found higher mean PRSs-SZ in capital and provincial areas compared to the other geographical areas. A PRS-SZ has been previously reported to be higher in people born in the capital area (Paksarian et al., 2018). Stratifying according to TRS status within geographical areas, we found a higher genetic liability for schizophrenia in people with TRS compared with those without TRS in the capital area. PRS-SZ levels were similar in people with or without TRS in the other geographical areas. Thus for people born in capital areas a 1 SD increase change in the PRS-SZ was associated with a 40% increased risk of TRS. In quartiles of the PRS, the risk for TRS was increased 2.4-fold in the highest quartile versus the lowest quartile of the PRS among people born in the capital area.

Another study from Denmark found significant correlation between a PRS-SZ with chronicity of schizophrenia as indicated by number and length of hospitalizations (Meier et al., 2016). As TRS is frequently detected only later in the course of treated schizophrenia, chronicity of the disorder may also play a role. As discussed by Gillespie and colleagues, “antipsychotics response and resistance are not readily separable from group differences in illness symptomatology or severity.” (Gillespie et al., 2017).

Beside polygenic risk for schizophrenia, a few other studies have documented a potential role of genetics in TRS. Ruderfer and colleagues recently reported genetic overlap, including genes represented by our applied polygenic risk score in this study, between schizophrenia pathogenesis and the antipsychotic mechanism of action (Ruderfer et al., 2016). Furthermore, the authors found rare disruptive variants in gene targets of antipsychotics and in genes with evidence for a role in antipsychotic efficacy in people with TRS (Ruderfer et al., 2016). Martin and Mowry found increased rare copy number variants in people with TRS (Martin and Mowry, 2016).

Our study has several limitations. First, we investigated the association between the potential risk factors urbanicity and PRS-SZ within the group of people with a registered diagnosis of schizophrenia. Our findings may be partly explained by this selection, i.e. collider-stratification bias (Cole et al., 2010). The collider-stratification bias may occur when two independent risk factors (here PRS-SZ and urbanicity) for schizophrenia are conditioned on their common effect (schizophrenia), which can impart an association between the two independent risk factors even though the association may not exist. This potential bias may complicate comparisons between our results and those obtained from other studies not conditioning on the status of schizophrenia in people with a diagnosis of TRS. Second, the predictive value of PRS-SZ based on the current known markers may be too unspecific to be of clinical utility, and other genetic markers of treatment resistant schizophrenia are needed. Third, the TRS definition was based on register information only, without clinical information on functioning or negative symptoms. We recently validated our algorithm against clinical records and found a PPV of 63.6%, and a sensitivity of 62.2% compared with a PPV of 78.3% and a sensitivity of 40% for identifying TRS based on clozapine prescriptions only (Ajnakina et al., 2018).

## 5 CONCLUSION

Among people with schizophrenia, we replicated previous findings of a weak genetic liability for schizophrenia and of place of birth in association with TRS. An interaction between urbanicity and PRS-SZ can be suspected with a PRS-SZ being predictive of TRS mainly in people with schizophrenia who resided in urban areas at birth.

## Acknowledgement

This study was funded by The Lundbeck Foundation, Denmark (Grant number: R155-2014-1724: Period II: 1 March 2015 – 28 February 2018).

This research has been conducted using the Danish National Biobank resource supported by the Novo Nordisk Foundation.

## Contributors

CG wrote the protocol and first draft of the manuscript, HTH analyzed the data. YW is responsible for genetic information and polygenic risk score for schizophrenia. All authors contributed to the design of the study. All authors contributed to and approved the final manuscript.

