## Supplemental Table 1 and Figure 1 for "Schizophrenia polygenic risk scores, urbanicity and treatment-resistant schizophrenia"

**Supplementary Table 1.** Cox regression analysis of **clozapine initiation** any time after schizophrenia diagnosis

| | No. of individuals | No. of events | Time at risk in years | Incidence rate per 100 person-years | HRs and 95% CIs for TRS | | |
|---|---|---|---|---|---|---|---|
| | | | | | Crude | Adjusted* | Adjusted** |
| **Polygenic risk score** | | | | | | | |
| Continuous | 4475 | 345 | 19 018 | 1.81 (1.63-2.02) | 1.01 (0.91-1.12) | 1.14 (0.99-1.30) | 1.14 (1.00-1.31) |
| Quartiles | | | | | | | |
| 0 (lowest) | 1118 | 78 | 4739 | 1.65 (1.32-2.05) | 1.00 (ref) | 1.00 (ref) | 1.00 (ref) |
| 1 | 1119 | 77 | 4739 | 1.62 (1.30-2.03) | 0.99 (0.72-1.35) | 0.99 (0.72-1.36) | 1.02 (0.74-1.39) |
| 2 | 1119 | 102 | 4717 | 2.16 (1.78-2.63) | 1.31 (0.98-1.76) | 1.35 (1.00-1.81) | 1.41 (1.05-1.90) |
| 3 (highest) | 1119 | 88 | 4822 | 1.83 (1.48-2.25) | 1.12 (0.82-1.52) | 1.31 (0.95-1.82) | 1.33 (0.96-1.86) |
| **Geographical area at birth** | | | | | | | |
| Capital | 1391 | 86 | 6121 | 1.41 (1.14-1.74) | 1.0 (ref) | | 1.0 (ref) |
| Provincial | 1702 | 145 | 7153 | 2.03 (1.72-2.39) | 1.41 (1.08-1.84) | | 1.38 (1.05-1.81) |
| Rural | 1382 | 114 | 5744 | 1.98 (1.65-2.38) | 1.38 (1.04-1.82) | | 1.36 (1.02-1.82) |

* Adjusted for 10 principal components.

** Different baseline hazards for males and females; mutually adjusted for polygenic risk score, age at diagnosis, year of diagnosis, geographical area at birth, type of genotyping array, 10 principal components.

IR: Incidence rate

HR: Hazard ratio

CI: confidence interval

**Supplementary Figure 1.** Mean polygenic risk score for schizophrenia according to geographical area at birth, adjusted for sex, age at diagnosis, year of diagnosis, geographical area at birth, type of genotyping array, 10 principal components.

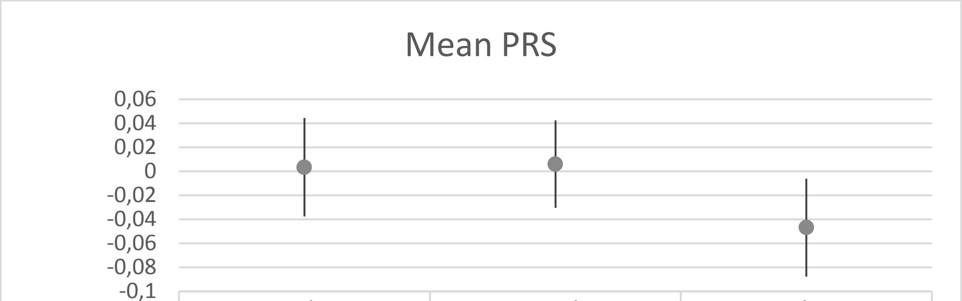

| | Capital area | Provincial area | Rural area |
|---|---|---|---|
| upper | 0,044427 | 0,042341 | -0,006029 |
| lower | -0,037726 | -0,03043 | -0,087612 |
| ● Mean PRS | 0,003351 | 0,005956 | -0,04682 |
